# Investigating the Effects of Osmolytes and Environmental pH on Bacterial Persisters

**DOI:** 10.1101/693366

**Authors:** Prashant Karki, Mehmet A. Orman

**Author notes:** Corresponding Author: S222 Engineering Bldg 1, 4726 Calhoun Rd, Houston, TX 77204.

## Abstract

Bacterial persisters are phenotypic variants that temporarily demonstrate an extraordinary tolerance towards antibiotics. Persisters have been linked to the recalcitrance of biofilm related infections; hence, a complete understanding of the physiology of persisters can lead to improvement of therapeutic strategies associated with such infections. Mechanisms pertaining to persister formation are known to be related to stress response pathways triggered from intra- or extra-cellular stress factors. Unfortunately, studies demonstrating the effects of osmolyte- and/or pH- induced stresses on bacterial persistence are largely missing. To fill this knowledge gap within the field, here we studied the effects of various osmolytes and pH conditions on *Escherichia coli* persistence with the use of phenotype microarrays and antibiotic tolerance assays. Although we found that a number of chemicals and pH environments, including urea, sodium nitrite and acidic pH, significantly reduced persister formation in *E. coli* compared to no-osmolyte/no-buffer controls, this reduction in persister levels was lessened in late-stationary-phase cultures. Our results further demonstrated a positive correlation between cell growth and persister formation, which challenges the general notion in the field that slow-growing cultures have more persister cells than fast-growing cultures.

## INTRODUCTION

Persisters are “a small fraction of a subpopulation in an isogenic bacterial culture that is temporarily tolerant to lethal concentrations of antibiotics”^1^. Persisters were first discovered by Gladys Hobby^2^, but the term was first coined by Joseph Bigger when he had encountered them in penicillin-treated *Staphylococcus pyogenes* cultures^3^. Unlike resistant bacteria, which undergo genetic mutations to avoid the action of antibiotics, persisters have the same genetic background as their antibiotic sensitive progenies^1,4^. After their discovery, persister cells had been largely ignored by the scientific community as their existence had not been considered an issue due to the emerging threat of antibiotic resistance. However, a groundbreaking study published by Balaban and colleagues^5^ reignited the interest in the field, encouraging many scientists to focus on various aspects of persistence phenomenon^5^.

Persister are considered to be an important health concern as their existence and ability to survive the antibiotic treatment has been linked to the recalcitrance of biofilm infections^1,6^. Persisters have been identified in every microorganism studied to date including highly pathogenic *Mycobacterium tuberculosis*^7^, *Staphylococcus aureus*^8^, *Pseudomonas aeruginosa*^9^ and *Candida albicans*^10^. Persisters also serve as a reservoir from which multi drug tolerant mutants can arise^11–12^. Substantial efforts in this field to molecularly characterize persistence have identified a number of mechanisms, which are generally associated with the SOS response^13–14^, the ppGpp-dependent stringent response^15^, reactive oxygen species (ROS)^16^, toxin/antitoxin (TA) modules^17^ and autophagy (i.e., self-digestion)^18^. Although stochastic dynamics of these cellular processes play a significant role in persister formation, environmental stresses such as nutrient depletion, hypoxia and overpopulation^19^ are also known to induce persistence. While the importance of carbon sources^20^, nutrients depletion^21^ and aerobic/anaerobic respiration^18,22^ to persister formation has been extensively studied by various research groups, the impacts of osmosis and environmental pH, which are known to be vital for cell growth and function, remain largely underexplored.

A focus on osmolytes as well as extra cellular pH could aid in identifying possible mechanisms related to persister formation and survival. Bacterial cells in principle need to maintain the osmotic pressure as it dictates the cell turgor which is considered to be a necessary force for cell growth and division^23^. Hence, it is essential for cells to adapt to any osmolarity changes occurring in their extracellular environment. Some of the molecules pertaining to osmoregulation, such as porins (e.g., OmpC and OmpF), solute specific efflux channels, mechanosensitive channels and osmoprotectant compounds, were already shown to be associated with microbial infections^24^. For instance, expression of *proP*, a gene encoding an osmosensing transporter, was found to be abnormally high in pathogenic *E. coli* isolates from urinary tract infection patients^25^. Similarly, mutation of *ompR*, a porin associated with osmoadaptation, was found to reduce virulence of *Shigella flexneri* and *Salmonella typhimurium*^26–27^. pH is another important factor that should be explored as recent studies demonstrated that acidic environments of macrophages (pH4-6.5) potentially induce persister formation^28^.

Our research goal in this study is to elucidate the effects of various osmolytes and pH on persister formation using a model organism, *Escherichia coli*, and Biolog MicroArray Plates (Biolog Inc., CA). Our analysis showed that although many osmolytes and pH>5 have minimum effect on *E. coli* persistence, we found a number of chemicals and pH conditions, such as urea, sodium nitrite and acidic environments (pH ≤ 5), reduced persister levels significantly. Our continuous effort further demonstrated that the observed reduction in persister levels was due to the delay in cell growth, resulted from osmolyte- and pH-induced stresses.

## MATERIAL AND METHODS

### Strain, Chemicals and Media

*E. coli* MG1655 used in this study was obtained from the Brynildsen Lab at Princeton University. Unless noted otherwise, all chemicals were purchased from Fisher Scientific (Pittsburg, PA). Ofloxacin (OFX) was purchased from VWR (Atlanta, GA). We used a modified Luria-Bertani (LB) medium for cell growth, composed of 1 g tryptone and 0.5 g of yeast extract in 100 ml of ultra-pure DI water and sterilized by autoclaving. Sodium Chloride (NaCl) was not added, in order to avoid any osmotic stress that is not arising from the osmolyte of interest. LB agar media (4 g premixed LB agar in 100 ml ultra-pure DI water, sterilized by autoclaving) were used to enumerate the colony forming units. Phenotypic arrays (PM-9 and PM-10 plates in 96-well formats) were obtained from Biolog Inc. (Hayward, CA) for screening the osmolytes and different pH conditions. PM-9 plates contain thirteen different osmolytes at various concentrations. PM-10 consists of various buffers and amino acids to control the culture pH. LB with 3.5% urea was prepared by dissolving 3.5 g urea crystals in 100 ml modified LB. LB with 60 mM sodium nitrite (NaNO_2_) was prepared by diluting 1 M NaNO_2_ in modified LB. LB with 100 mM 2-N-morpholino-ethanesulfonic acid (MES) at pH 5 was prepared by mixing 50 ml of 2x LB and 10 ml of 1 M MES pH 5 solution with ultra-pure DI water to achieve a final volume of 100 ml. LB with MES pH 7; 3-morpholinopropane-1-sulfonic acid (MOPS) pH 7; and MOPS pH 8 media were prepared using the same protocol by replacing the MES pH 5 content with corresponding buffer solution. The 2x LB medium was prepared by adding 2 g of Tryptone and 1 g of Yeast extract in 100 ml ultra-pure DI water and sterilized by autoclaving. M-9 Glucose media were prepared by mixing 10 ml 100 mM Glucose solution, 20 ml 5XM-9 salt, 0.2 ml of 1 M Magnesium Sulphate (MgSO_4_) and 0.01 ml of 1 M Calcium Chloride (CaCl_2_) with ultra-pure DI water to achieve a final volume of 100 ml^20^. For persister selection 5 mg/ml of OFX stock solution was prepared by dissolving 5 mg of OFX salt in 1 ml of ultra-pure DI water with 10 μl of 10M Sodium hydroxide (NaOH) solution. NaOH was added to fully dissolve OFX in water. For persister assays, 5 μg/ml OFX was used^18,20,22^. MIC ranges for *E. coli* MG1655 were found to be 0.039-0.078 μg /ml for OFX by using a method based on serial 2-fold dilutions of antibiotics in 2 ml LB media in 14-ml test tubes^29^. All solutions and media were filter sterilized to avoid any contamination.

### Growth Conditions

Overnight pre-cultures were prepared from a 25% glycerol cell stock (−80°C) in 2 ml modified LB media (without salt) in 14-ml test tubes and cultured at 37°C with shaking (250 rpm) for 12 h. Then, the cultures were diluted at 1:100 in 25 ml of fresh modified LB in 250-ml baffled flasks. The cultures’ optical densities were monitored at 600 nm (OD_600_) using Varoskian Lux plate reader (ThermoFisher, Waltham, MA). Upon reaching an OD_600_ value of ~ 0.5, the cells were diluted at 1:100 in 250-ml baffled flasks containing fresh, modified LB and cultured to achieve OD_600_ ~ 0.5. This dilution/growth cycle was performed to minimize the carryover of persisters from the overnight cultures^30^. After propagating the cell cultures twice, cells were diluted 100 fold in fresh, modified LB one more time, and 150 μl cell suspensions were then transferred to each well in the PM plates (Fig. 1A). Controls that do not have osmolytes or buffers were prepared by transferring the cell suspensions to empty 96-well plates. Half-area flat-bottom plates were used to be consistent with the area of the PM plates. The plates were covered with sterile, oxygen-permeable sealing membranes and cultured at 37°C and 250 rpm. The pre-propagated cells from the dilution/growth cycle experiments were also inoculated (1:100) into 25 ml LB media with different osmolytes (i.e., urea, NaNO_2_) or pH in 250-ml baffled flasks. After culturing the cells for 24 h, persister assays were conducted as described below.

**Fig. 1:**
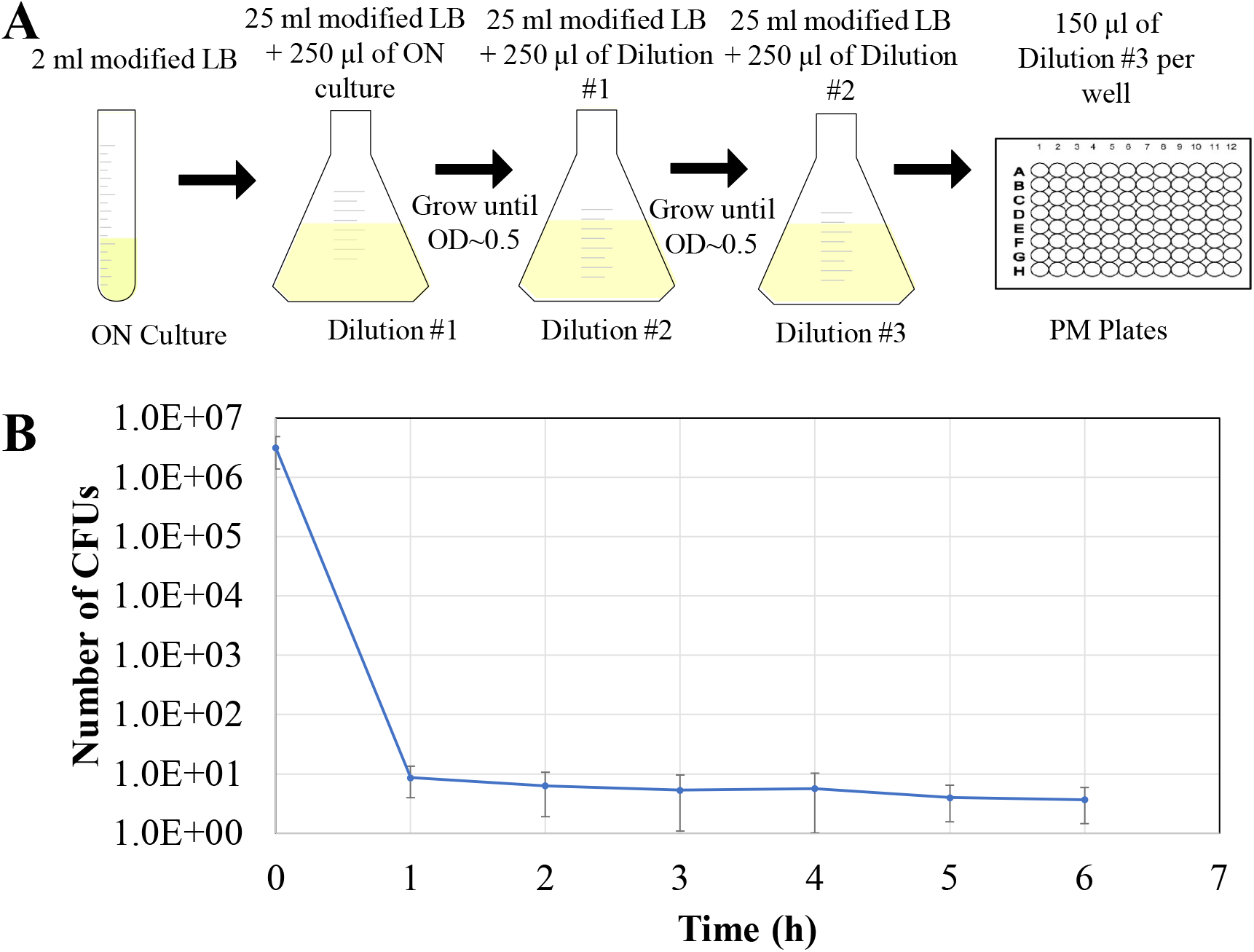
Eliminating pre-existing persisters in ON cultures. (A) Overnight cultures were diluted 1:100 fold in modified LB and grown until mid-exponential phase for the dilution/growth cycle experiments. After the second cycle, the cells were diluted 1:100 fold in fresh modified LB transferred to microarray plates. (B) The cells in the third dilution flask were treated with 5 μg/ml OFX. At designated time points, 1 ml samples were collected, washed to remove the antibiotic, serially diluted and spotted in LB agar plate to count the number of survived cells as measured by CFUs.

### PM-Plate Screening

After 24 h of growth, 10 μl cell suspensions were removed from each well of PM plates and transferred (20-fold dilution) to regular flat-bottom 96-well plates where each well has 190 μl modified LB with 5 μg/ml OFX. After 6 h-antibiotic treatment, 10 μl samples from each well were serially diluted in 290 μl phosphate-buffered saline (PBS)/well in round-bottom 96-well plates. Then, 10 μl diluted samples were spotted on LB agar plates and incubated for 16 h at 37°C to measure colony forming units (CFUs)^18^. As the LB cell suspension was transferred to PBS solution without washing off the antibiotic, the cells in PBS are exposed to an OFX concentration ≤0.16 μg/ml. However, we have shown that this concentration range was negligible enough to not alter the cell growth on the agar plates where OFX should be further diluted (Supp. Fig. 1).

### Testing the Identified Osmolytes and pH Conditions in Flasks

The pre-propagated cells after dilution/growth cycle experiments were inoculated (1:100) into 25 ml LB media with different osmolytes or buffers in 250-ml baffled flasks: Urea (3-4%), NaNO_2_ (60 mM-100mm), MES pH 5, MES pH 7, MOPS pH 7, MOPS pH 8 and controls (no- osmolyte and no-buffer). After culturing the cells in the presence of these chemicals for 24 h, cells were inoculated (1:20 or 1:100) in 25 ml fresh modified LB with 5 μg/ ml OFX in 250-ml baffled flasks. At designated time points, 1 ml samples from antibiotic-treated cultures were collected, washed with PBS solution to dilute the antibiotic to sub-MIC levels and resuspended in 100 μl of PBS. Ten μl cell suspensions in PBS were then serially diluted, plated on LB agar media and incubated for 16 h at 37°C to quantify the persisters.

### Growth State Dependence of Persistence

To assess the dependence of persister formation on cell growth in the aforementioned cultures, we adopted a method developed by Keren *et al.*^30^. Briefly, the pre-propagated cells after dilution/growth cycle experiments were inoculated (1:100) in 25 ml LB media with different osmolytes or buffers in 250-ml baffled flasks as described above. At designated time points, 500 μl cell cultures from the flasks were collected and mixed with 500μl of fresh LB with osmolytes or buffers in 14-ml test tubes, and then treated with 5 μg/ml of OFX for 6 h. We have already shown that 6-h treatment was sufficient to achieve biphasic kill curves (Fig. 1B). Cells before and after the treatments were washed to remove the antibiotic and plated on LB agar media to enumerate persisters.

To properly evaluate the impact of these pre-propagation cycles on persistence, these cycles were also performed in the modified LB with osmolytes and buffers. Overnight cultures were inoculated (1:100) into 25 ml of LB with the indicated osmolytes or buffers and cultured until OD_600_~0.5. After repeating the dilution/growth cycle twice in the presence of chemicals, the cell suspensions were inoculated 1:100 in 25 ml fresh LB with the corresponding chemicals. The persister levels throughout cell growth in the third set of flasks were determined using the method of Keren *et al.*^30^, as described above.

### Cell Growth and Persistence in Regular LB or M-9 Glucose Media

Overnight cultures were inoculated (1:100) in 25 ml regular LB with 1% NaCl or M-9 media with 10 mM glucose in 250-ml baffled flasks and grown at 37°C with shaking at 250 rpm At designated time points, persister levels were determined using the method of Keren *et al.*^30^, as described above.

### Quantifying Cell Growth and Persister Formation Rates

We calculated the cell growth and persister-formation rates at exponential phase by using a classical cell growth equation: *N*_*i*_ = *N*_0_ ∗ exp(*r* ∗ *t*) , where N_o_ is initial number of cells, *r* is the rate of growth or formation, *t* is the age of the culture and *N*_*i*_ is the number of cells at time “*i*”. The rates were determined using an Excel function, SOLVER, by minimizing the residual sum of squared differences (SSE) between the experimental data and the predicted data for exponential-phase cultures.

### Statistical Test

Three biological replicates were performed for each experimental condition. Each data point represents the mean value and the error bars represent the standard deviations of the data sets. Statistical significance was evaluated using the two-tailed t-tests with unequal variance as persister datasets were shown to be normally distributed^20^. The threshold for significance was set to be P<0.05. ANOVA test was used to statistically assess the correlation between cell growth vs. persister formation.

## RESULTS

Using the Biolog Phenotype (PM) plates, we aimed to study the effects of osmolytes and pH conditions on *E. coli* MG1655 persistence. We used a modified LB broth (without NaCl) in cell cultures to avoid any additional osmotic effect that could arise. Studies have shown that absence of NaCl (1%) from the regular LB broth does not alter the cell viability or cell-growth rates significantly^31^, which was also verified here (Supp. Fig. 3). For antibiotic tolerance assays, we chose OFX, a widely used quinolone, which is known to effectively kill both fast- and slow-growing cells^1,32^. Given that persisters are largely formed by passage through stationary phase, the persisters carried over from overnight cultures may hinder the effects of osmolytes and pH on persister formation in PM plates. Therefore, we used a strategy developed by Keren *et al.*^30^ to eliminate these phenotypic variants in cell cultures before transferring them to PM plates. Briefly, cells from overnight cultures were diluted (1:100) in fresh media, cultured until they reached exponential phase, and then diluted in fresh media again (Fig. 1A). As verified by Karen *et al.*^30^ and our results (Supp. Fig. 7), persisters formation is little or negligible at lag and early exponential phase. Hence, repeating the dilution/growth cycle twice in this growth phase reduced persister levels to the limit of detection (Fig. 1B). After the second cycle, cells were transferred to PM plates and grown for 24 h before treating them with OFX for persister quantitation. Cells were plated for CFU enumuration before and after the OFX treatments to assess the effects of osmolytes/pHs on *E. coli* cell viability and persistence, respectively.

### Sodium Nitrite, Urea and Acidic pH Reduce Persister Survival

Through PM-9 plates, we screened for sodium chloride, potassium chloride, sodium sulphate, ethylene glycol, sodium formate, urea, sodium lactate, sodium phosphate, sodium benzoate, ammonium sulphate, sodium nitrite and sodium nitrate at indicated concentrations (Figs. 2 and Supp. Fig. 2). Urea (4%) (Fig. 2B) and NaNO_2_ (80 mM) (Fig. 2C) reduced persister levels below the limit of detection while posing a minimal effect on cell viability, as measured by CFU levels before the OFX treatment. Although sodium chloride is known to be essential component of cell culture media, the concentrations lower than 8% did not significantly impact the *E. coli* cell viability and persistence; however, higher concentrations significantly compromised the cell viability (Figure 2A). Most of the chemicals tested either led to compromised cell viability at high concentrations (e.g., sodium formate and sodium benzoate) or had no net effect on persister levels within a wide range of concentrations tested (e.g., sodium sulfate, ethylene glycol, potassium chloride and sodium nitrate) (Supp. Fig. 2B, C, D, F, H, I, J and K). There were some cases (sodium lactate and ammonium sulphate) where persister levels were reduced; however, either the errors associated with the results were high or there was no noticeable trend observed across the concentrations studied (Supp. Fig. 2E and G). Further, a number of chemicals, including trigonelline, trimethylamine and trehalose, exhibited antipersister activities without affecting the *E. coli* cell viability in the presence of sodium chloride (Supp. Fig. 2A).

**Fig. 2:**
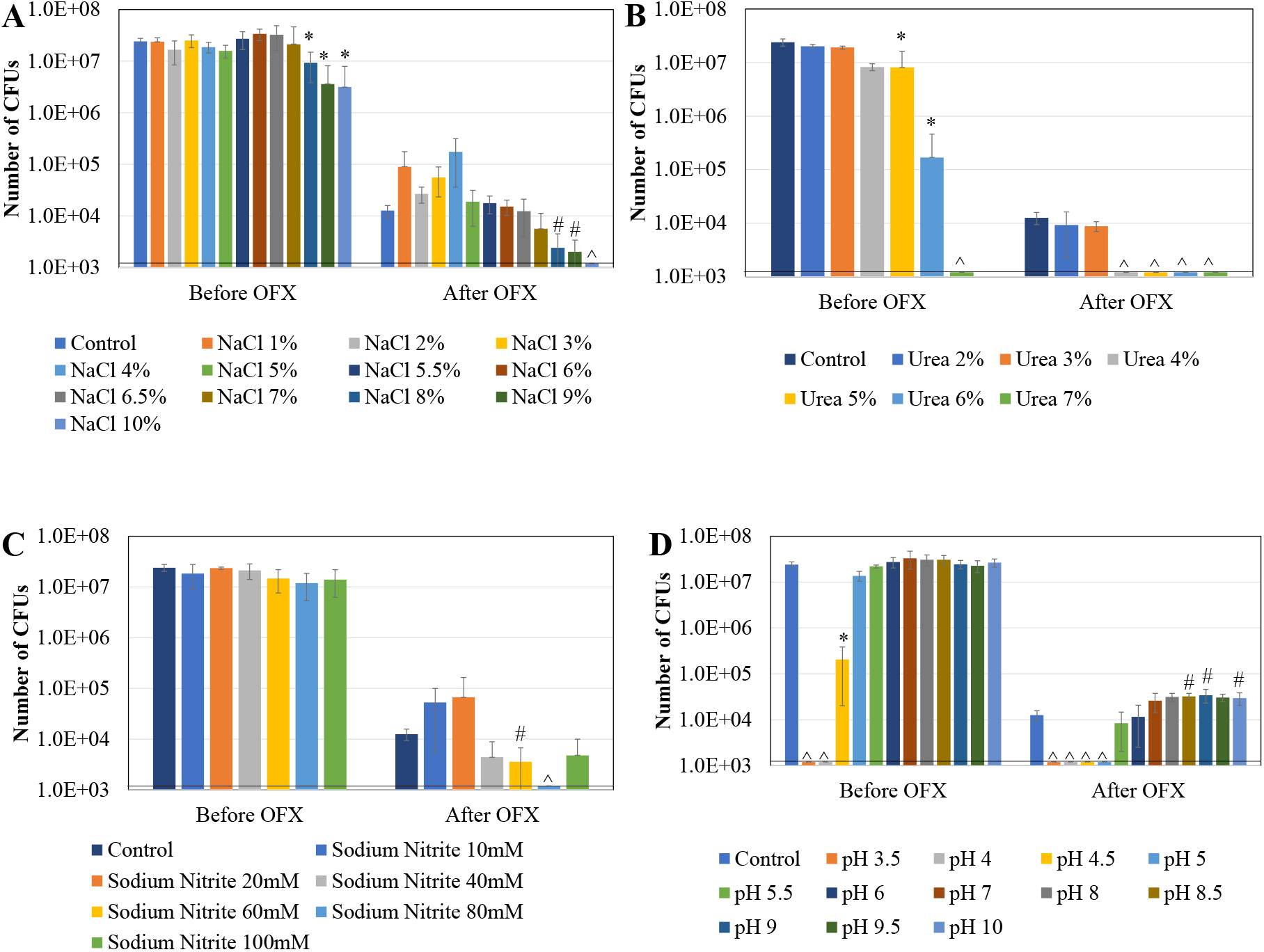
Persister levels in PM plates. Pre-propagated cells after the dilution/growth cycle experiments were transferred to PM plates and cultured for 24 h. Then, the cells from the plates were transferred to fresh media and treated with 5 μg/ml of OFX for 6 h. CFU measurements were performed before and after the OFX treatments. The effects of sodium chloride, urea, sodium nitrite and various pH conditions on E. *coli* cell survival and persistence are shown in A, B, C, and D panels, respectively. * indicates that the osmolyte or pH of interest significantly affects the *E. coli* cell viability compared to no-osmolyte or no-buffer controls before the antibiotic treatments (P<0.05). # indicates that the osmolyte or pH of interest significantly affects the OFX persister levels compared to no-osmolyte or no-buffer controls (P<0.05). ^ indicates CFUs below the limit of detection.

PM 10 plate contained various pH conditions (Fig. 2D). In contrast to our expectation, persister levels of *E. coli* in low pH environment were observed to be less than those in untreated control group while higher pH resulted in an increase or minimal effect in persister levels (Fig. 2D). However, at pH below 4.5, the *E. coli* cell viability was compromised and not many cells survived the OFX selection pressure. In addition to pH changes, we also tested various amino acids at low or high pH environments to study the effects of decarboxylation or deamination of amino acids on bacterial persistence (Supp. Fig. 2L and M). At low pH (4-5), *E. coli* cells are capable of producing decarboxylase enzymes that can produce amines from amino acids, thus, increasing the pH of the environment^33^. On the contrary, at alkaline pH, cells could have active enzymes that lead to deamination of amino acids^33^. Although addition of amino acids at indicated pH did not affect persistence mostly, lysine and glycine at pH 4.5 as well as homoserine at pH 9 significantly altered persister levels, without affecting the *E. coli* cell viability, compared to no-amino acid controls (Supp. Fig. 2L and M). Decarboxylation of lysine produces cadaverine which has been shown to improve cell survival in acidic environment^34^. Cadaverine induces closure of porin channel^35–36^ which could lower the transport of antibiotic across the cell membranes, leading to higher antibiotic tolerance. Homoserine, on the other hand, is an intermediate for glycine, serine and threonine metabolism. Although the effects of homoserine or its products on persisters has not been investigated, a study has shown L-homoserine, when converted to alpha-amino-n-butyric acid, led to inhibition of *Mycobacterium tuberculosis* growth^37^. Whether these effects on persistence were due to the buffering capabilities of amino acids or products of their reactions warrants further investigation.

### Reduction in Persister Levels are not Influenced by Culture Conditions

Persister levels are sensitive to culture and antibiotic-treatment conditions, including media type, culturing time, media volume, flask type, aeration and growth stage. To test whether the observed reduction in persister levels from PM cultures is reproducible, we recreated similar culture conditions using 250-ml baffled flasks. After propagating cells in the modified LB with the dilution/growth cycle experiments as described above (Fig. 1A), cells were transferred (1:100 dilution) to flasks containing modified LB with various osmolytes and buffers (e.g., urea, NaNO_2_, or buffers for low pH) that lead to a significant reduction in persistence in PM plates. After culturing the cells for 24 h in the presence of selected chemicals (Supp. Fig. 3), the cells were diluted (1:100) in fresh modified LB with OFX in flasks for persister enumeration (Fig. 3). We first tested the effects of osmolytes of interest at various concentrations (Supp. Fig. 4A and 4B), and found that 3-4% urea and 60-100 mM NaNO_2_ significantly reduced the persister levels compared to no-osmolyte controls, consistent with the PM results (Fig. 2). When we tested various buffers (MES pH 5, MES pH 7, MOPS pH 7 and MOPS pH 8) to test the effects of pH, we found persister levels in MES pH 5 media were significantly lower compared to no-buffer controls while MOPS pH 8 did not affect the *E. coli* persistence (Supp. Fig. 4C). The similar persister levels in MES pH 7 as well MOPS pH 7 conditions (Supp. Fig. 4C) validate that the observed reduction in persistence was related to the pH of the environment not the buffer salts. We note that persister assays here were performed after diluting cells 100-fold in fresh, modified LB with OFX in flasks. We also wanted to test the effects of cell density in assay cultures as it has been related to quorum sensing in bacteria^38,39^. However, the observations from two different inoculation rates (1:20 vs. 1:100) used for persister assay cultures followed the similar trends (Fig. 3 vs. Supp. Fig. 5). Overall, these results obtained in baffled flasks at higher volumes allowed us to confirm that the reported persister levels were due to the osmolytes and the varying environmental pH, not resulted from culture conditions such as aeration or volume.

**Fig. 3:**
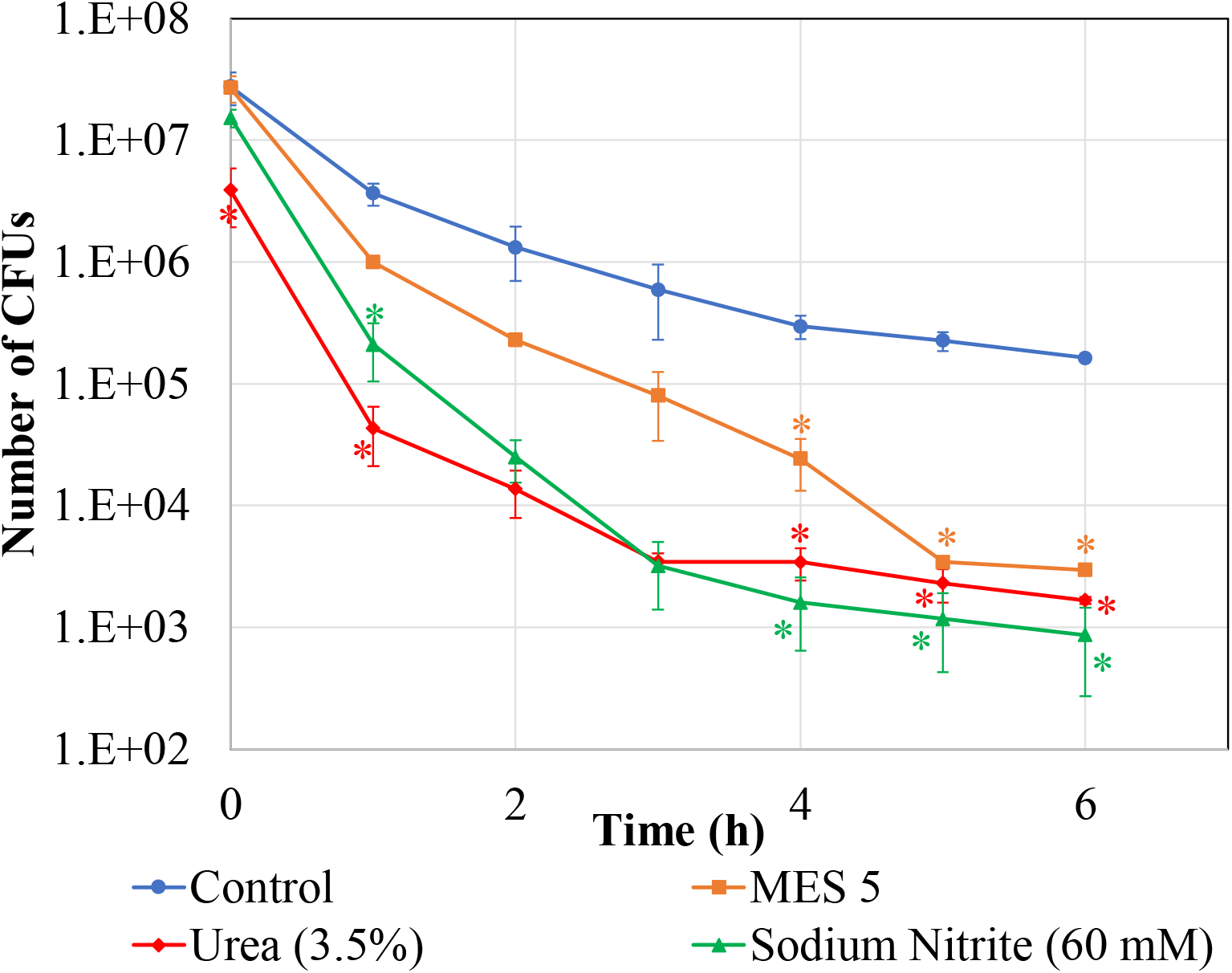
Effects of Urea (3.5%), NaNO_2_ (60 mM) and pH 5 on *E. coli* persistence. Pre-propagated cells were diluted 1:100 fold in fresh, modified LB media with urea, NaNO_2_ and MES 5 at indicated concentrations in baffled flasks. After growing the cultures for 24 h, the cells were diluted (1:100) to fresh, modified LB media with 5 μg/ml of OFX. At designated time points, samples were collected, washed to remove the antibiotics and plates on agar media to quantify CFUs. * indicates the statistical difference between the treatment and control groups (P<0.05).

### Persister Formation Positively Correlates with Cell Growth

Cells grown in media containing urea, NaNO_2_ or MES pH 5, entered the stationary phase at later time points compared to no-osmolyte and no-buffer control groups (Supp. Fig. 3). To test if this delay influenced the rate at which persisters were being formed in the cultures, we adopted a method developed by Keren *et al.*^30^. Briefly, at indicated time points (Fig. 4), 500 μl of the samples from the cultures were collected and mixed with 500 μl fresh media (including an osmolyte or buffer) and treated with OFX to quantify persisters. We observed that cells cultured in the presence of osmolytes or buffers grew slower and formed persisters at slower rates compared to control groups (Fig. 4). The rate of persister cell formation seems to correlate positively with the rate of cell growth at exponential-growth phase in these cultures. In addition, the observed differences between persister levels were lessened at late-stationary phase (t = 36 h).

**Fig. 4:**
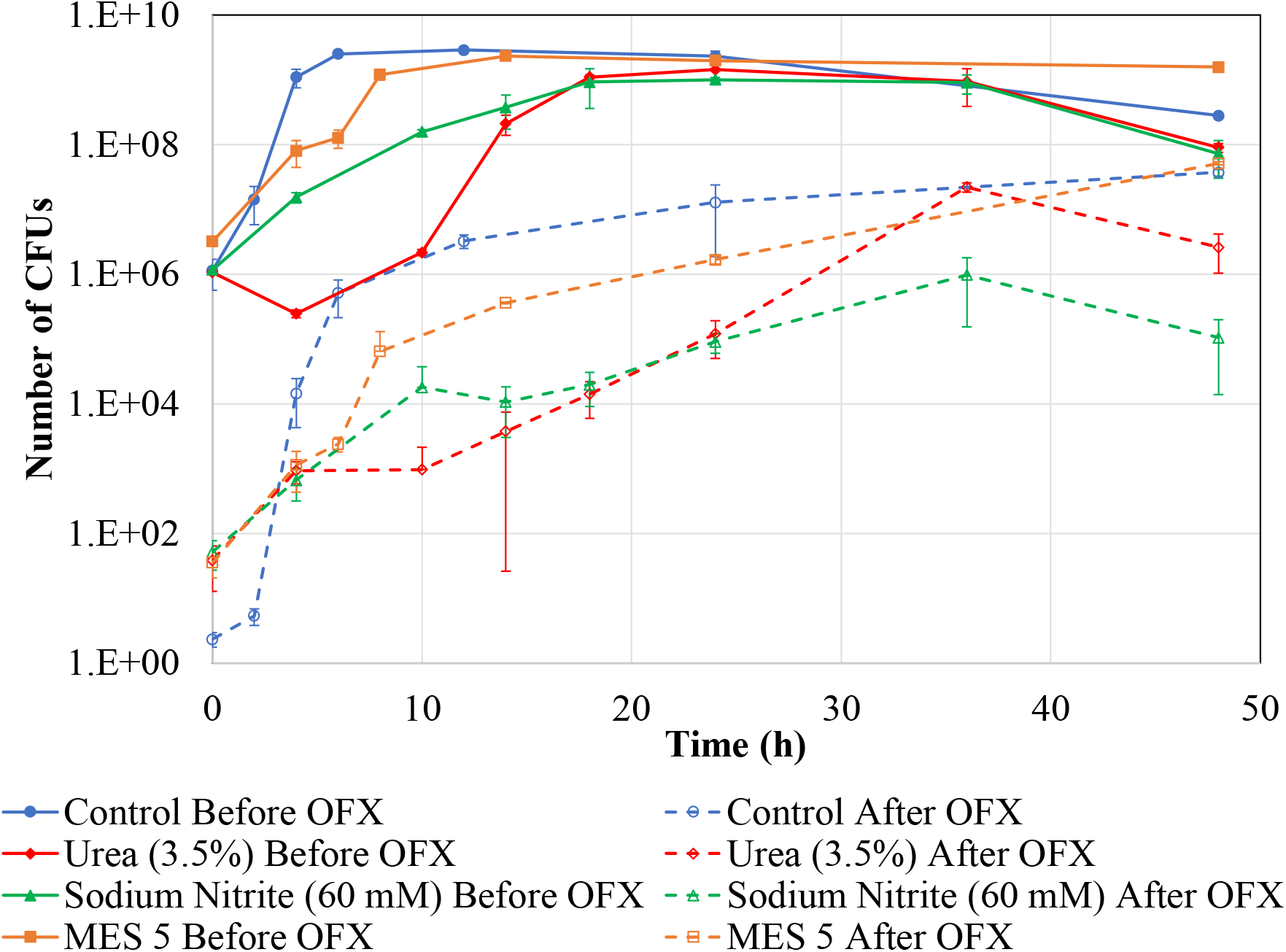
Dependence of persister formation on cell-growth. Pre-propagated cells were transferred at 1:100 ratio in modified LB containing osmolytes and pH buffers as indicated in baffled flasks, and cultured for 48 h. At designated time points, samples were collected and diluted 1:2 in fresh media with osmolytes and buffers and treated with 5 μg/ml OFX for 6 h. The solid lines indicate CFUs measured before antibiotic treatments and the dashed lines indicate CFUs measured after the antibiotic stress (persisters).

Transition of cells from dilution/growth cycles (Fig. 1A) to new environments can temporarily inhibit the cell growth, thus affecting persistence. When we performed these dilution/growth cycles in the presence of osmolytes or buffers, we observed the same growth patterns in all conditions except the cultures containing urea (Supp. Fig. 6). Upon growing the cells continuously in LB with urea, the extended lag phase generated by the bacteriostatic property of urea was not observed (Fig. 4 vs. Supp. Fig. 6). Like the results shown in Fig. 4, persister levels in late-stationary-phase cultures (t=36 h) were similar across all conditions studied (Supp. Fig. 6). The positive correlation between the cell growth and persister formation patterns at exponential-growth phase still exists in both experimental setups (Fig. 4 and Supp. Fig. 6). To statistically analyze this correlation, we calculated exponential-phase cell growth and persister formation rates in these cultures using a classical cell growth equation (see Materials and Methods) and generated a “cell growth vs. persister formation” plot (Fig. 5). Using ANOVA with the null hypothesis that the two variables are not linearly related, we verified that the observed positive correlation is statistically significant (P < 10^−15^). To show if this trend is ubiquitously observed in various environmental conditions and not specific to osmolyte- and pH-induced stresses, cells from overnight cultures were diluted 1:100 in flasks containing regular LB (fast-growing culture media) or M-9 Glucose (slow-growing culture media) and cultured to monitor their growth and persistence without performing the dilution/growth cycles. As expected, cell growth and persister formation rates were higher in LB compared to those in M-9 Glucose (Supp. Fig. 7), further supporting the proposed correlation (Fig. 5). This interesting observation challenges the current paradigm that slow growing cell cultures potentially have more persister cells than fast-growing cultures.

**Fig. 5:**
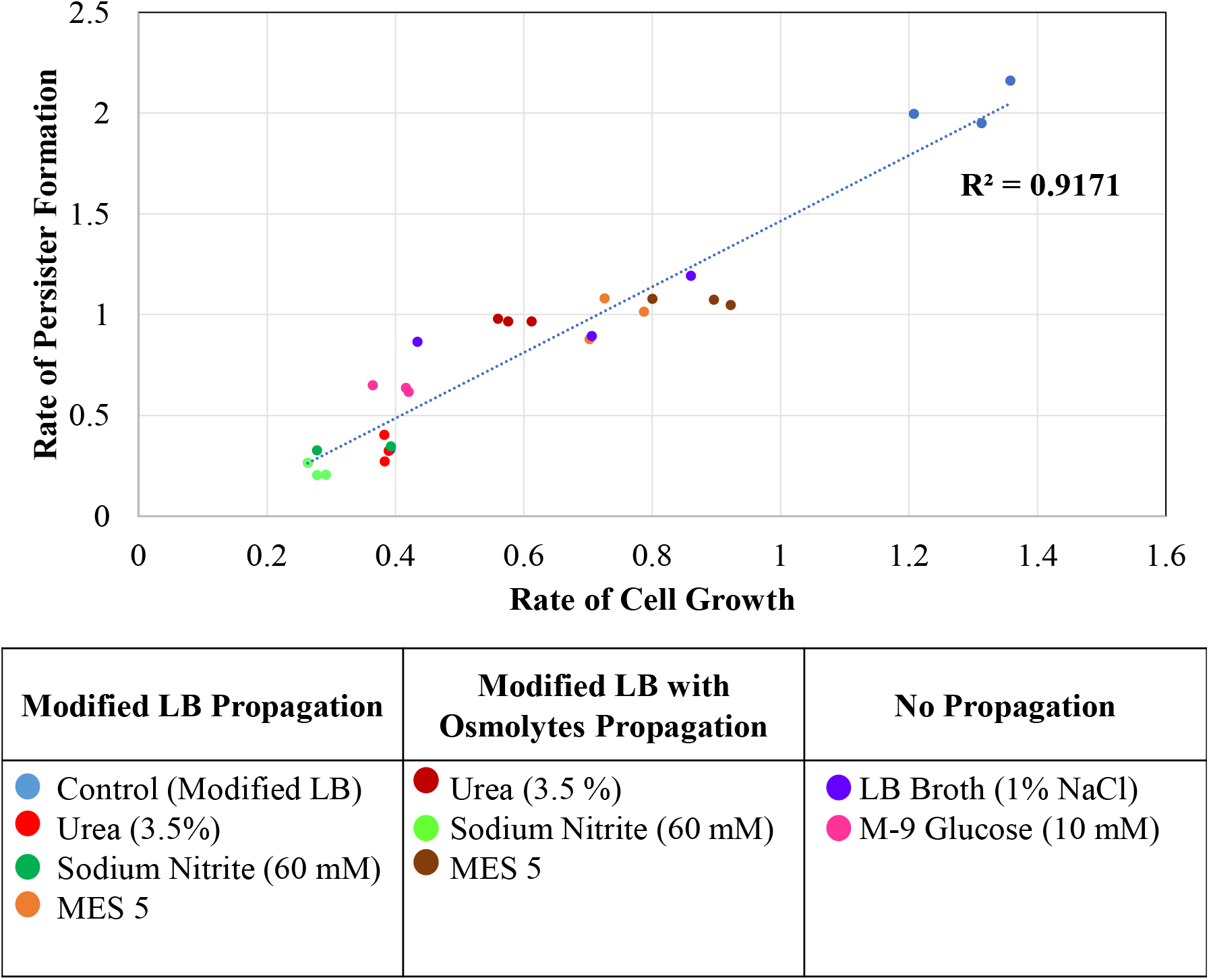
Correlation between the cell growth and persister formation at exponential growth phase. The rates were calculated using the traditional cell growth equation (*N*_*i*_ = *N*_*0*_ ∗ exp(*r* ∗ *t*)) for all conditions indicated.

## DISCUSSION

Osmosis is an integral part of cell functions; osmotic pressure impacts cell growth rate and turgor as well as transport phenomena between the cell and the extracellular environment^23^. Here, we studied the impacts of various osmolytes on *E. coli* persistence by screening a number of chemicals with various concentrations in PM plates. For most of these compounds at extreme conditions (e.g., NaCl>8% or pH<4), the *E. coli* cell viability was significantly compromised. On the other hand, a number of chemicals, including urea and sodium nitrite, significantly reduced persister levels at certain concentrations while slightly hampering the viability of the *E. coli* cells (Fig. 3).

Urea has been shown to possess antimicrobial properties that have been under investigation for a while^40,41^. Bacteria undergo pleomorphism and have abnormal morphological forms in the presence of urea^42^. We anticipated that the change in osmolarity of the environment with urea may have led to hypertonicity in the cells; however, such was not found to be true according to previous studies^40^. Although the underlying mechanism is not well understood, urea can act as a bacteriostatic at 3.5 % and higher concentrations in growth media^40^, similar to our study where we observed little to no cell growth in the first 8 hours of cell cultures (Fig. 4 and Supp. Fig. 3). Sodium nitrite, which is used as a food preservative, was also shown to have bacteriostatic effects^43–44^. This has been linked to inhibition of glucose transport, oxygen uptake and oxidative phosphorylation^45^. These factors are potential contributors to the slow growth leading to a prolonged lag as observed in this study. In addition, inhibition of respiration has been shown to reduce degradation of endogenous proteins and ribosomal RNAs in stationary-phase cultures, thus increasing the susceptibility of stationary-phase cells to antibiotics when transferred to fresh media^18^. Whether such mechanisms exist in cells grown in the presence of urea or sodium nitrite should be investigated further.

Along with osmolytes, pH is another factor that may affect bacterial persistence. Physiological pH in various regions of host body varies from acidic (e.g., stomach linings, macrophages) to basic pH (e.g., liver)^46,47,48^. Although we generally did not observe a significant difference in persister levels for pHs > 5, highly acidic environment (pH ≤ 5) reduced persister cells in *E. coli* cultures (Fig. 2). Helaine *et al.* showed that pre-exposing the *Salmonella* cells to acidic LB (pH 4.5) for 30 min significantly increased persister formation^28^. Kim Lewis’ group, on the other hand, showed that culturing the cells in low pH environment for 2-3.5 h did not significantly alter *E. coli* persistence although they observed an increase in the expression of *dinJ/yafQ*, *mqsRA*, *relBE and yefM/yoeB* TA systems^49^. We note that the assay conditions studied by these groups are considerably different than those studied here. Culturing time of cells under a stress condition seems to be an important factor (Fig. 4), which might explain this reported disparity.

In our studies we have shown that slow cell growth mostly results in slow rate of persister formation, unlike the general notion in the field. Although such dependence of persisters on the growth stage of cell cultures has been shown and verified in a few studies^5,30,50^, these studies were mostly conducted in media without any added environmental stimuli. We expect that growth related molecules, such as ppGpp, reactive-oxygen species (ROS) or quorum sensing molecules (e.g., indole), could explain the positive correlation between cell growth and persistence since these molecules are known to result in a phenotypic switch from a normal, growing state to a non-growing, persistence state in bacteria^51^. Cell ageing may also play a role here as faster growing cells age at a more rapid pace leading to a higher rate of protein aggregation, which have been shown to be correlated with persistence^51^.

Through this screening study, we have generated a considerable amount of data which will significantly contribute to the existing persister knowledgebase, given that greater knowledge of persister physiology will aid development of novel anti-persister strategies. These findings may be used to design a co-treatment strategy (e.g., antimicrobial ointments) as the identified chemicals (e.g., sodium nitrite, trigonelline, trimethylamine and trehalose) can serve as adjuvants to enhance antibiotic therapies against persistent infections (e.g., biofilm related wound infections). In addition to clinical significance, the growth dependent nature of persister formation further verifies that culture conditions, the time of antibiotic treatments and the age of cultures are important factors that should be taken into consideration in persister assays.

## ACKNOWLEDGEMENTS

We would like to thank Sayed G. Mohiuddin for providing assistance in persister assays and the members of Orman Lab research group for their valuable contributions to this projects. The research was supported by NIH/NIAID K22AI125468 Career Transition award and University of Houston start up grant.

