## Supplementary File for "Investigating the Effects of Osmolytes and Environmental pH on Bacterial Persisters"

**Department of Chemical and Biomolecular Engineering, University of Houston,**

**Houston, TX, 77204**

* Corresponding Author
S222 Engineering Bldg 1
4726 Calhoun Rd
Houston, TX 77204


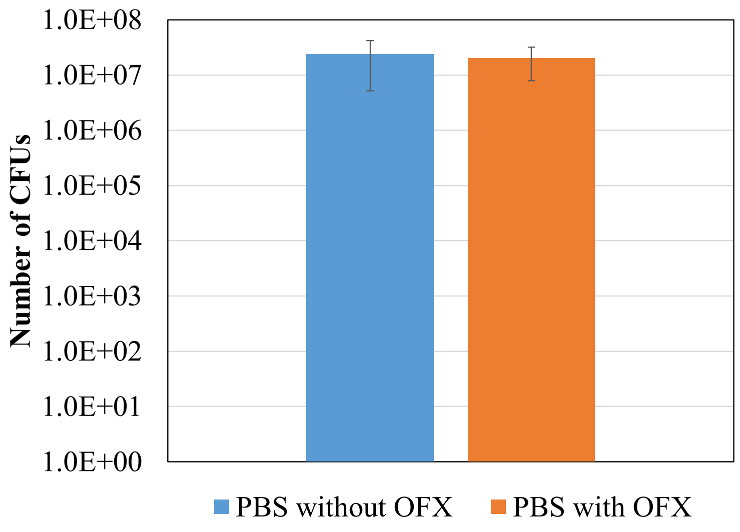


**Supp. Fig. 1: Comparison of cell viability in OFX concentration in unwashed cell suspension.** Overnight cultures were diluted 1:100 in LB media. Cells were then grown for 4 h at 37°C shaking at 250 rpm. At 4 h, 10 µl cell cultures were serially diluted (10-fold) six times in 90 µl PBS. Similarly, 10 µl cell cultures were also serially diluted in 90 µl PBS containing 0.166 µg/ml of OFX. Then, diluted samples were plated on LB agar plates.

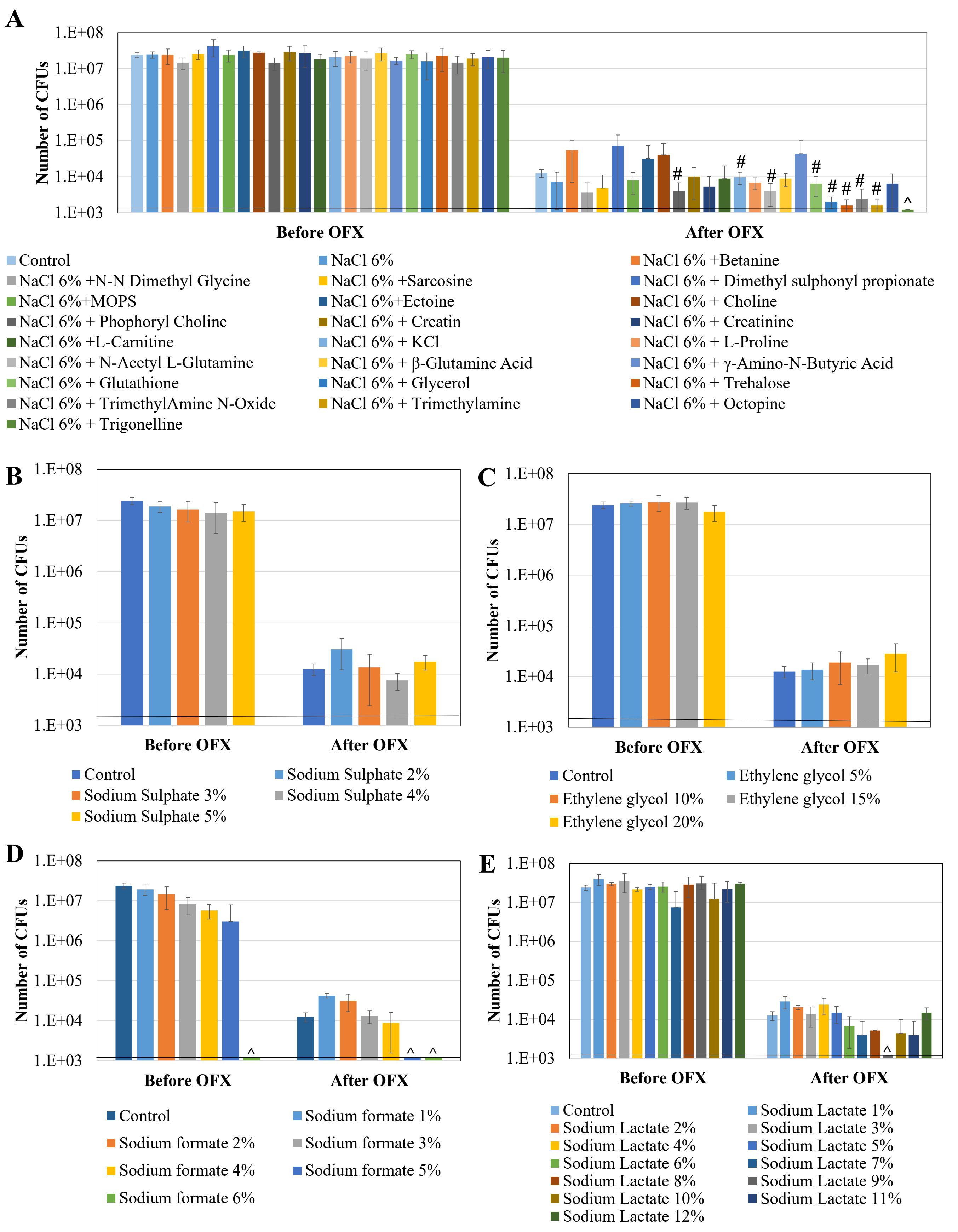


**Supp. Fig. 2: Continued below**

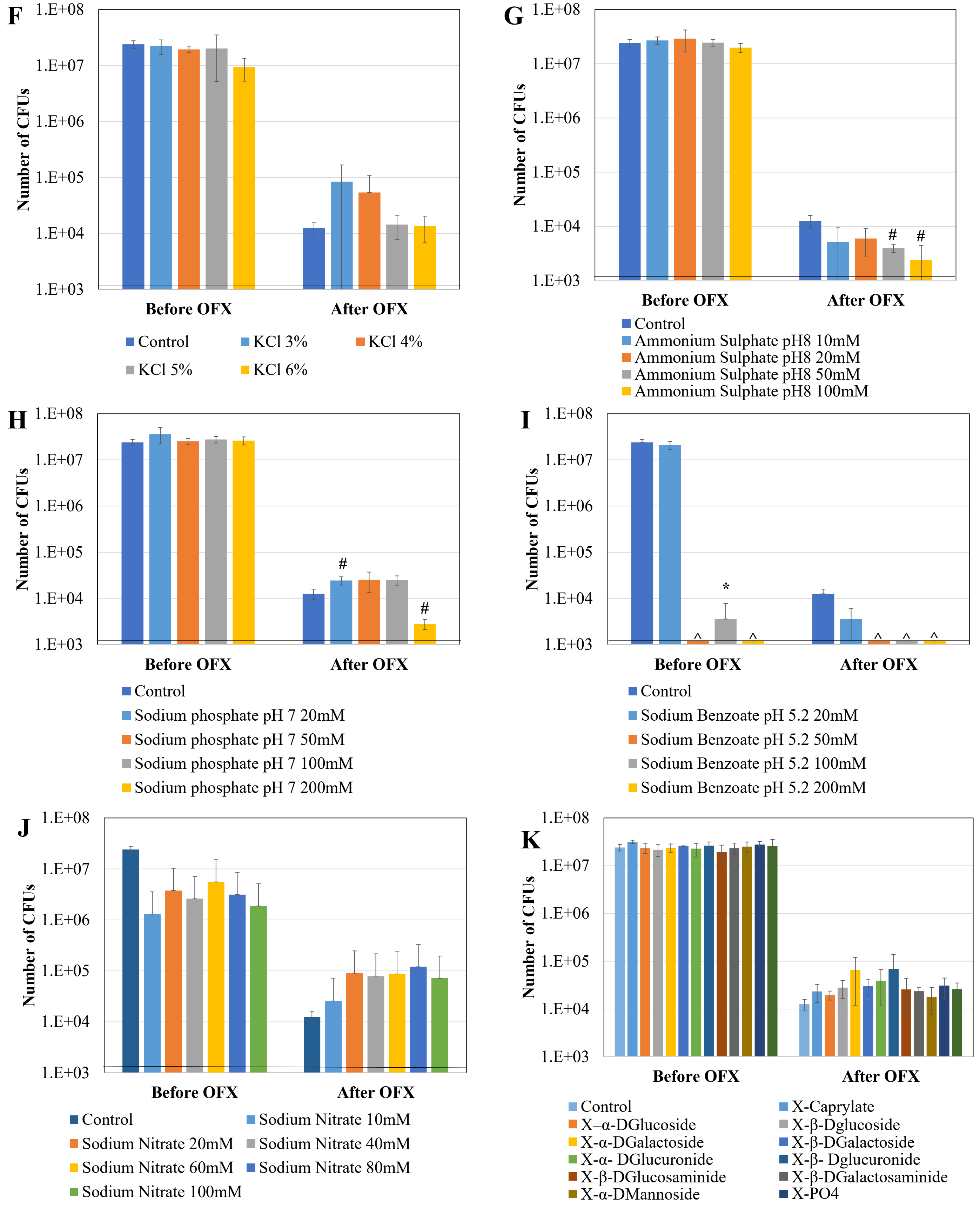


**Supp. Fig. 2: Continued below**

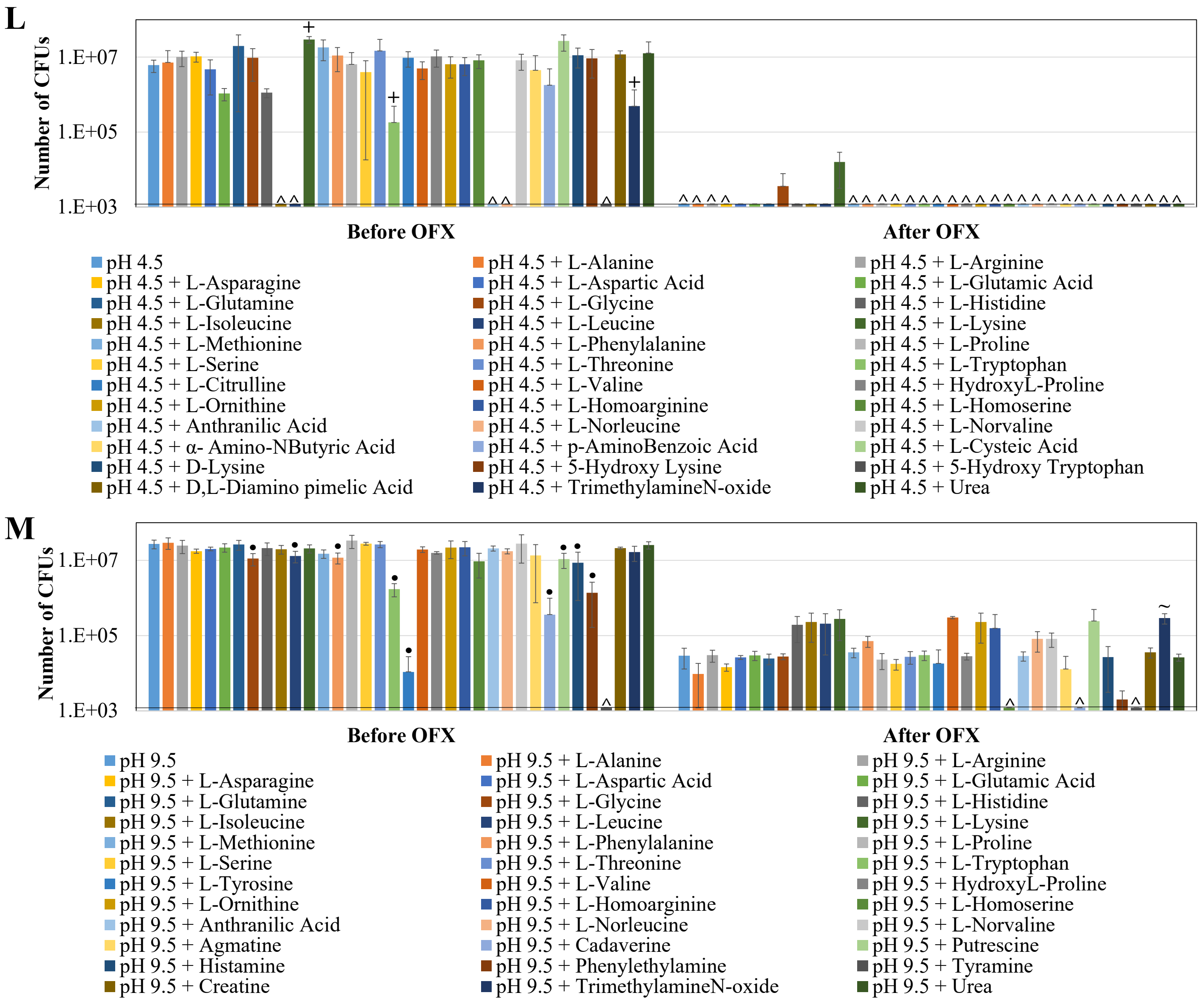


**Supp. Fig 2: Persister levels in PM plates.** Pre-propagated cells after the dilution/growth cycle were transferred to PM plates and cultured for 24 h. Then, the cells from the plates were transferred to fresh media and treated with 5 µg/ml of OFX for 6 h. CFU measurements were performed before and after the OFX treatments. *,+ and • indicate that the osmolyte or pH of interest significantly affects the *E. coli* cell viability compared to “no-osmolyte”, “no-amino acid at pH 4.5” or “no-amino acid at pH 9.5” controls, respectively, before the antibiotic treatments (P<0.05). # and ~ indicate that the osmolyte or pH of interest significantly affects the OFX persister levels compared to “no-osmolyte” or “no-amino acid at pH 9.5” controls respectively (P<0.05). ^ indicates CFUs below the limit of detection.

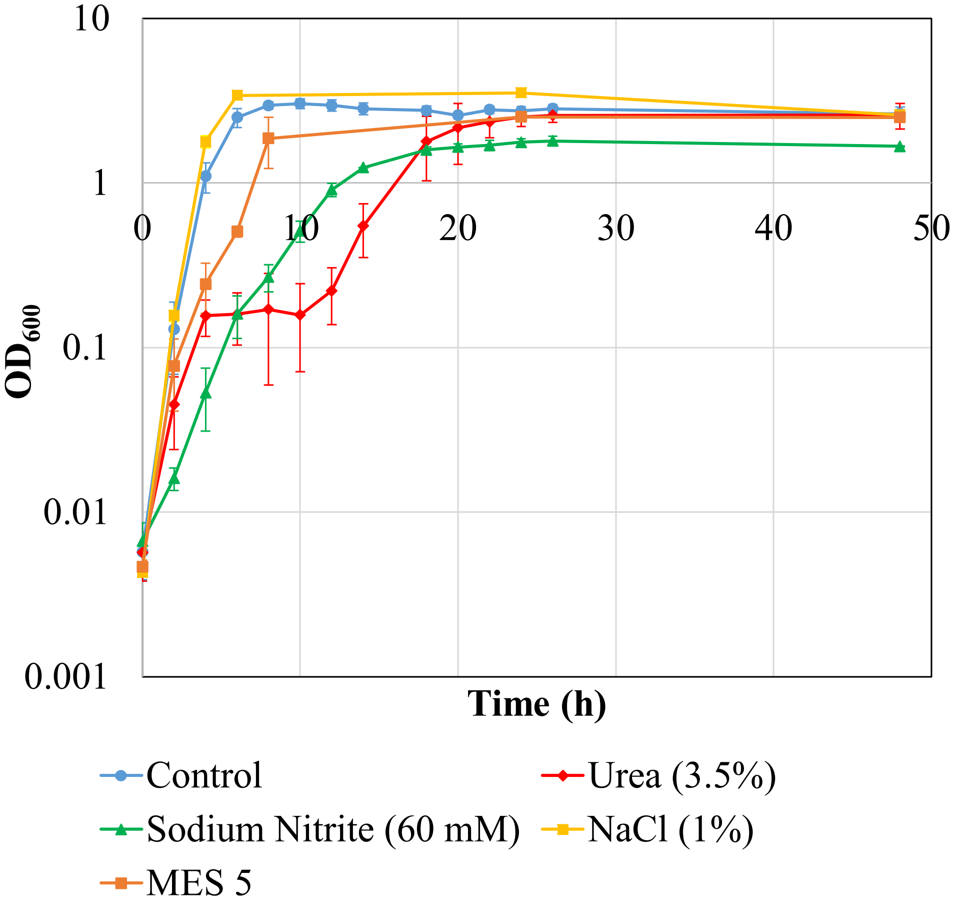


**Supp. Fig 3: Growth curves of cell cultures containing osmolytes and buffers.** Pre-propagated cells were transferred at 1:100 ratio in modified LB containing osmolytes (1 % NaCl, 3.5% Urea and 60 mM $NaNO_2$) and pH (5) buffers in baffled flasks, and cultured for 48 h. At designated time points, samples were collected to measure $OD_{600}$.

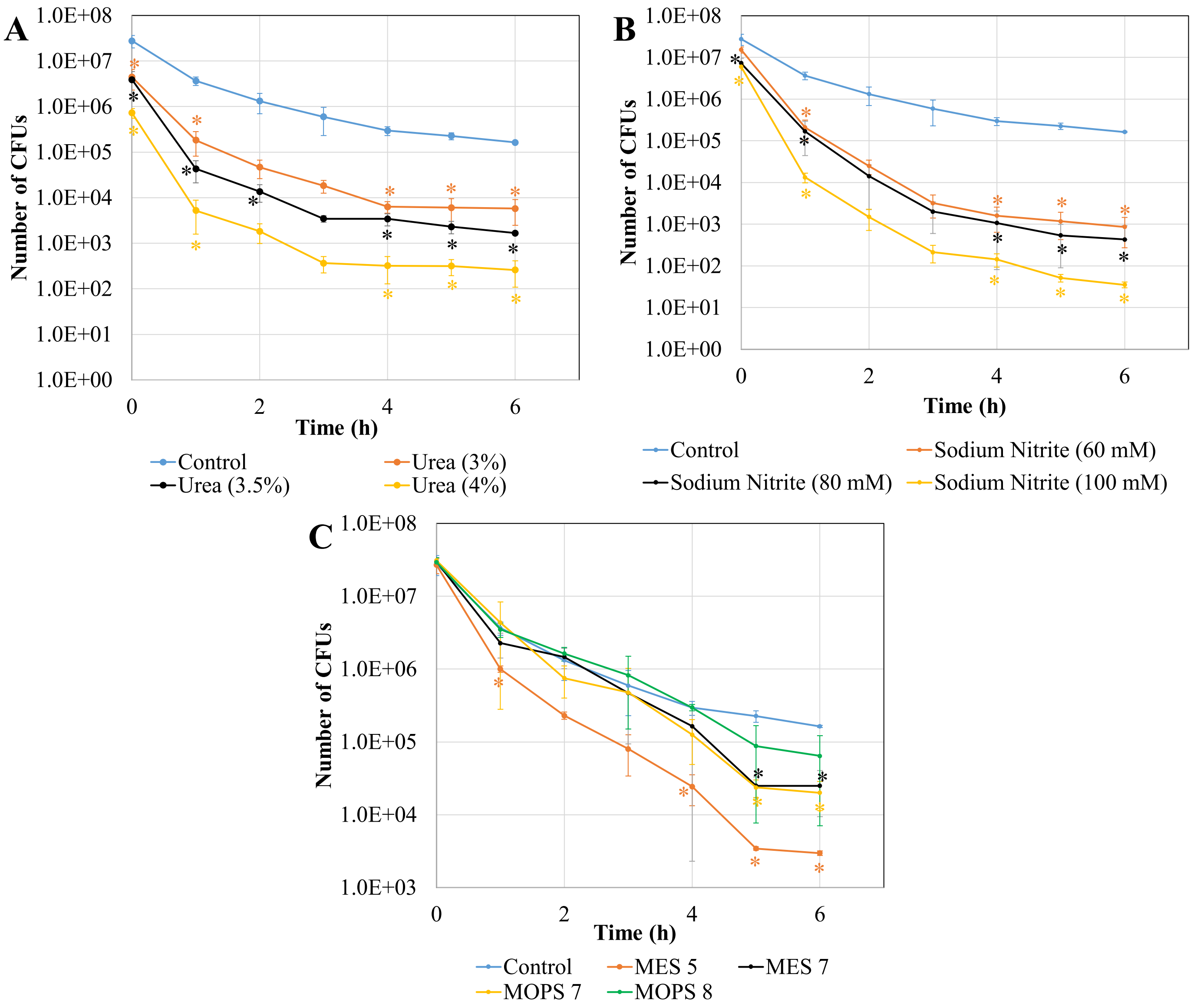


**Supp. Fig. 4: Effects of osmolytes and buffers on *E. coli* persistence.** Pre-propagated cells were diluted 1:100 fold in fresh, modified LB media with indicated osmolytes and pH buffers in baffled flasks. After growing the cultures for 24 h, the cells were diluted (1:100) to fresh, modified LB media with 5 µg/ml of OFX. At designated time points, samples were collected, washed to remove the antibiotics and plates on agar media to quantify CFUs. * indicates the statistical difference between the treatment and control groups ($P<0.05$).

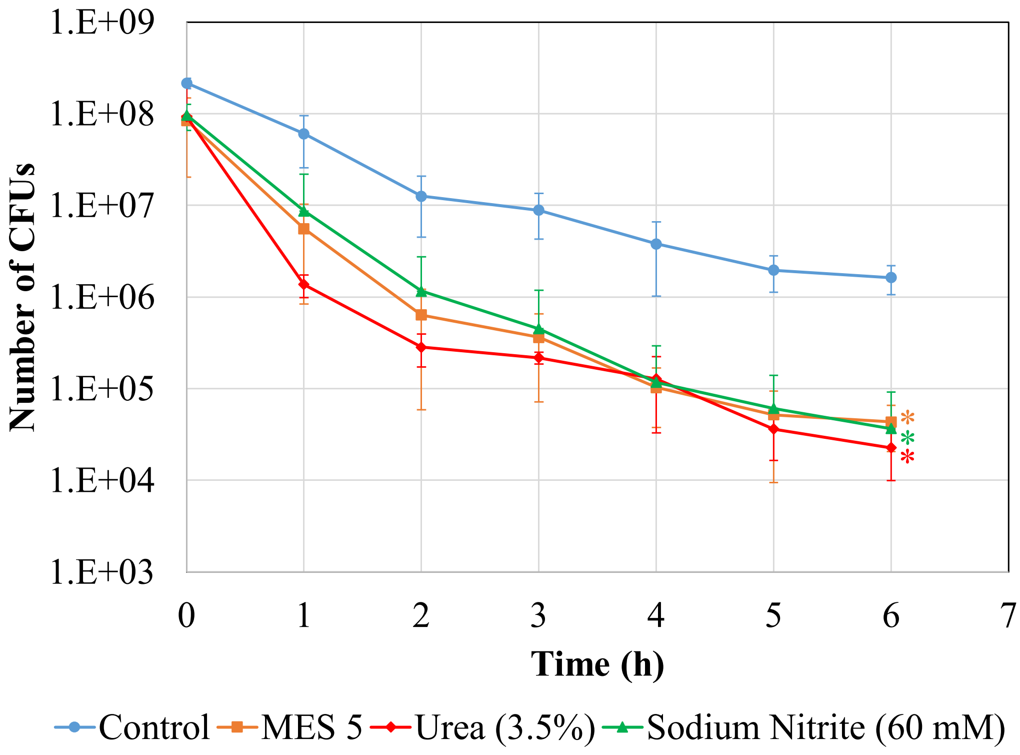


**Supp. Fig. 5: Effect of inoculation rate on *E. coli* persistence**. Pre-propagated cells were diluted 1:100 fold in fresh, modified LB media with indicated osmolytes (1 % NaCl, 3.5% Urea and 60 mM NaNO2) and pH buffers in baffled flasks. After growing the cultures for 24 h, the cells were diluted (1:20) to fresh, modified LB media with 5 µg/ml of OFX. At designated time points, samples were collected, washed to remove the antibiotics and plates on agar media to quantify CFUs. * indicates the statistical difference between the treatment and control groups ($P<0.05$).

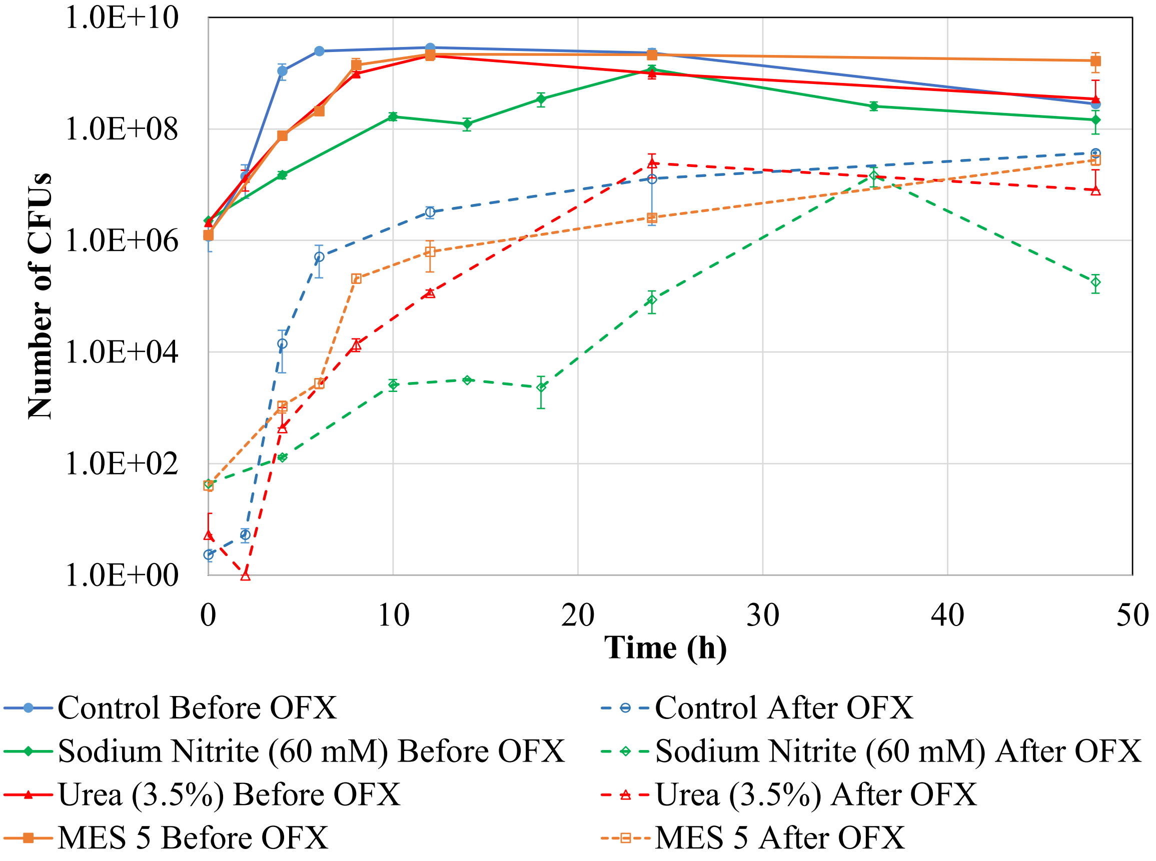


**Supp. Fig. 6: Dependence of persister formation on cell-growth.** Overnight cultures were diluted 1:100 in modified LB containing osmolytes (3.5% Urea and 60 mM $NaNO_2$) and pH buffers, grown until mid-exponential phase. This dilution/growth cycle repeated once more in the presence of osmolytes and buffers. Then, cells were then transferred at 1:100 ratio in modified LB containing osmolytes and pH buffers and cultured for 48 h. At designated time points, samples were collected and diluted 1:2 in fresh media with osmolytes and buffers and treated with 5 μg/ml OFX for 6 h. The solid lines represent CFUs measured before antibiotic treatments and the dashed lines represent CFUs measured after the antibiotic stress (persisters).

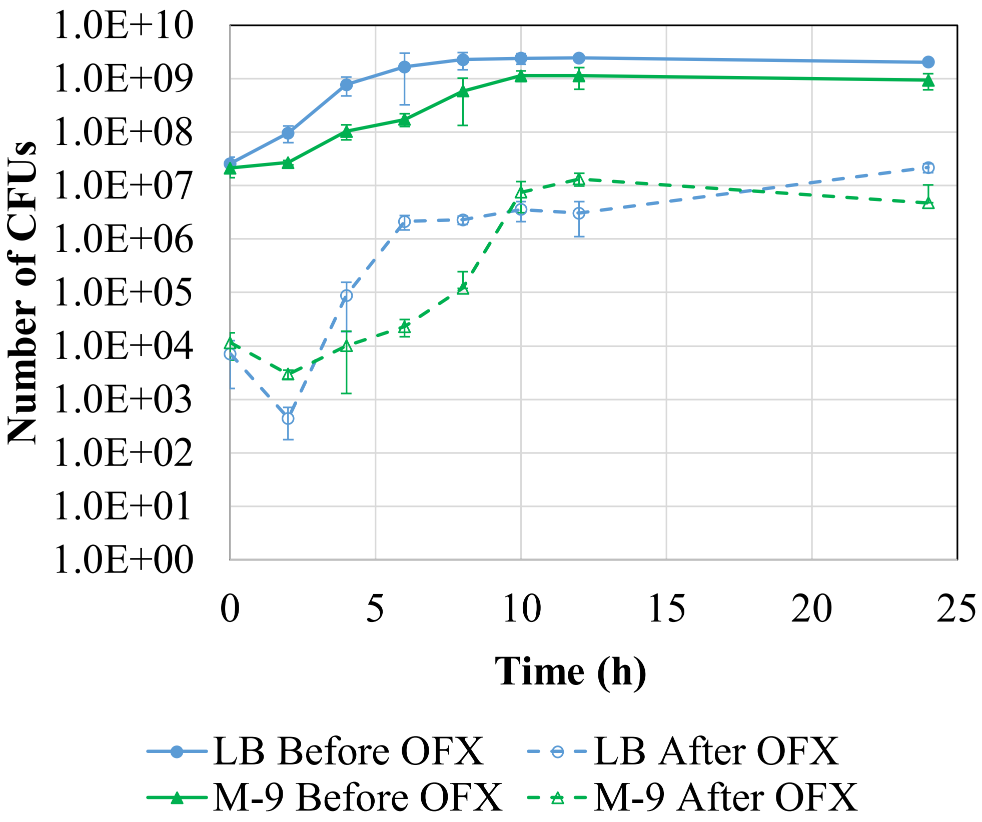


**Supp. Fig. 7: Persister formation in cultures grown in LB Broth and M-9 Glucose media.** Overnight cultures were diluted 1:100 in regular LB and M-9 Glucose media at 37°C while shaking at 250 rpm for 24 h. At designated time points, samples were collected and diluted 1:2 in their corresponding media and treated with 5 µg/ml OFX for 6 h. The solid lines represent CFUs measured before antibiotic treatments and the dashed lines represent CFUs measured after the antibiotic stress (persisters).
